# Interleukin-8/CXCR2 signaling regulates therapy-induced plasticity and enhances tumorigenicity in glioblastoma

**DOI:** 10.1101/454553

**Authors:** Tanwir Hasan, Seamus P. Caragher, Jack M. Shireman, Cheol H. Park, Fatemeh Atashi, Shivani Baisiwala, Gina Lee, Donna Guo, Meijing Wu, Jennifer Y. Wang, Mahua Dey, Maciej S. Lesniak, Craig M Horbinski, C. David James, Atique U. Ahmed

**Author notes:** Correspondence; The Department of Neurological Surgery, Feinberg School of Medicine, Northwestern University, 303 East Superior Street Chicago, IL 60611. These authors contributed equally (Tanwir Hasan, Seamus P. Caragher).

## Abstract

Glioblastoma (GBM) remains one of the least treatable types of cancer. Recent work highlights two key factors contributing to this resistant phenotype—cellular plasticity, the ability of GBM cells to adopt many phenotypes, and the microenvironment. Here, we provide evidence that Interleukin-8 (IL-8) plays a vital role in promoting cellular plasticity and cancer initiating cells (CICs) niche during anti-glioma chemotherapy. IL-8 expression is significantly elevated during chemotherapy, and immunohistochemical analysis of matched primary and recurrent patient GBM tissues revealed about 60% of recurrent tissues IL-8 expression is upregulated. In silico analysis of the TCGA data indicated that IL-8 signaling could promote epigenetic plasticity by altering the polycomb repressor complex activity. We are proposing that such regulation my promote epigenetic plasticity, which allows the GBM cells to adapt therapy and may promote therapeutic resistance. Our data show that IL-8 is a crucial microenvironmental factor involved in developing therapeutic adaptation and can be targeted in combination with conventional chemo-and radiotherapy to prevent disease recurrence.

**ABSTRACT:** Emerging evidence reveals enrichment of glioma initiating cells (GICs) following therapeutic intervention. This enrichment occurs partly by dedifferentiation of non-GICs to GICs within the tumor, which may contribute to therapeutic resistance and the generation of lethal recurrent tumors. To elucidate the molecular mechanisms governing therapy-induced cellular plasticity, we performed genome-wide chromatin immunoprecipitation sequencing (ChIP-Seq) and gene expression analysis (gene microarray analysis) during treatment with standard of care temozolomide (TMZ) chemotherapy. Analysis revealed significant enhancement of open chromatin marks in known astrocytic enhancers for Interleukin-8 (IL-8) loci as well as elevated expression during anti-glioma chemotherapy. The Cancer Genome Atlas and Ivy Glioblastoma Atlas Project data demonstrated that IL-8 transcript expression is negatively correlated with GBM patient survival (p=0.001) and positively correlated with that of genes associated with the CIC phenotype such as KLF4, c-Myc and HIF2α (p<0.001). Immunohistochemical analysis of patient samples demonstrated elevated IL-8 expression in about 60% of recurrent GBM tumors relative to matched primary tumors and this expression also positively correlates with time to recurrence. Exposure to IL-8 significantly enhanced the self-renewing capacity of patient-derived xenograft (PDX) GBM (average 3-fold, p<0.0005). Furthermore, IL-8 knockdown significantly delayed PDX GBM tumor growth *in vivo* (p<0.0005). Finally, guided by *in silico* analysis of TCGA data, we examined the effect of therapy-induced IL-8 expression on the epigenomic landscape of GBM cells and observed increased trimethylation of H3K9 and H3K27. Our results show that IL-8 alters cellular plasticity and mediates alterations in histone status. These finding suggest that IL-8 signaling participates in regulating GBM adaptation to therapeutic stress and therefore represents a promising target for combination with conventional chemotherapy in order to limit GBM recurrence.

## INTRODUCTION

Glioblastoma (GBM) is the most aggressive and prevalent primary brain tumor in adults, with 10,000 new diagnoses each year. Recurrent tumors, with increased invasive and resistance capacities, are an inevitability for GBM patients despite aggressive therapeutic intervention. Glioma-initiating cells (GICs) are considered a key driver of primary tumor development, as well as major contributors to tumor recurrence [1].

Recent reports demonstrate that differentiated GBM cells undergo cellular and molecular changes to acquire GIC-like states[2, 3]. This cellular plasticity dramatically complicates our ability to prevent tumor recurrence in the clinical setting. Our group and others have shown that therapeutic stress and microenvironmental dynamics induce cellular plasticity in GBM, driving the conversion of differentiated GBM cells to the GIC states [3–5]. However, the exact mechanisms governing post-therapy GBM plasticity remain unknown. Unraveling the signaling pathways that drive this plasticity will provide key insight for improving treatment of GBM.

Using gene expression and bioinformatics analysis, we identified Interleukin-8 (IL-8) as a key player in promoting post-therapy cellular plasticity. IL-8 enhances the selfrenewal capacity of patient-derived xenograft (PDX) GBM cells and is elevated in recurrent GBM patient specimens. Reducing levels of IL-8 in murine models significantly improved survival and enhanced the efficacy of Temozolomide chemotherapy. We demonstrate that IL-8/CXCR2 signaling alters the epigenomic landscape in GBM cells, inducing a GIC-like state and increasing the proportion of GICs after treatment. This study indicates that IL-8 signaling influences GBM plasticity and recurrence, highlighting it as a potential novel therapeutic target in GBM.

## Materials and Methods

### Cell Culture

U251 human glioma cell lines were procured from the American Type Culture Collection (Manassas, VA, USA). These cells were cultured in Dulbecco’s Modified Eagle’s Medium (DMEM; HyClone, Thermo Fisher Scienftific; San Jose, CA, USA) supplemented with 10% fetal bovine serum (FBS; Atlanta Biologicals, Lawrenceville, GA, USA) and 1% penicillin-streptomycin (P/S) antibiotic mixture (Cellgro; Herdon, VA, USA; Mediatech, Herdon, VA, USA). Patient-derived xenograft (PDX) glioma specimens (GBM43, GBM12, GBM6, GBM5, and GBM39) were obtained from Dr. C. David James at Northwestern University and maintained according to published protocols [6].

### Animals

Athymic nude mice (nu/nu; Charles River; Skokie, IL, USA) were housed according to all Institutional Animal Care and Use Committee (IACUC) guidelines and in compliance with all applicable federal and state statutes governing the use of animals for biomedical research. Briefly, animals were housed in shoebox cages with no more than 5 mice per cage in a temperature and humidity-controlled room. Food and water were available *ad libitum*. A strict 12-hour light-dark cycle was maintained.

### Co-immunoprecipitation

For co-immunoprecipitation (Co-IP) experiments, proteins were extracted and quantified as described above. Then, 50-100μg of protein were incubated with primary antibody overnight at 4°C with gentle rocking. The next day, anti-rabbit IgG antibodies conjugated to agarose beads were added to the cell lysates and incubated for at least 1 hour at 22°C. Next, the mixture was spun down and washed several times in PBS. Finally, proteins were eluted from the mixture and loaded into gels, as described above.

### Human Sample Histology

Human primary and matched recurrent GBM tissue were obtained from the Northwestern University’s Nervous System Tumor Bank. All patients were consented according to Institutional Review Board (IRB) policies prior to the obtainment of samples. Samples were formalin-fixed and paraffin-embedded (FFPE). Immunohistochemistry of tumor samples was performed on 4-μm-thick sections heated at 60 °C for at least 1 hour. Staining for IL-8 was carried out manually and antigen retrieval was performed with a Biocare Medical Decloaking Chamber using high (LC3) or low pH antigen retrieval buffer from Dako. Primary antibodies were incubated for 1 h at room temperature. A secondary antibody was EnVision labeled polymer-HRP (horseradish peroxidase) anti-mouse or antirabbit as appropriate. Staining was visualized using 3, 3’-diaminobenzidine (DAB) chromogen (Dako, K8000).

IL-8 immunohistochemical results on TMAs were semiquantified on a relative scale from 0 to 3, with 0 = negative and 3 = strongest (see Supplemental Figure 1). Each tumor was represented by 3 separate cores on 3 separate blocks.

### Statistical analysis

All statistical analyses were performed using the GraphPad Prism Software v4.0 (GraphPad Software, San Diego, CA, USA). Where applicable, one-way ANOVA, unpaired t-test, and log-rank test were applied. Survival distributions were estimated with the Kaplan–Meier method. A P value <0.05 was considered statistically significant.

For detailed methods and material please see the Supplementary Methods section.

## Results

### Therapeutic stress increases IL-8 expression in vitro and in vivo

To investigate if Temozolomide (TMZ) chemotherapy promotes the adoption of a GIC state via cellular plasticity, gene set enrichment analysis (GSEA) using the Affymetrix platform and genome-wide ChIP-Seq analysis were performed. Data from patient-derived xenograft (PDX) GBM43 cells 4 and 8 days post-treatment with either vehicle control (DMSO) or physiological doses of TMZ (50μM) [7–9]revealed a significant (FDR q = 0.08, FWER p-value = 0.046) enrichment of a network of genes responsible for supporting the GIC phenotype (Figure 1A)[10]. In order to identify potential novel regulators of this process, we examined a variety of other genes for enhanced expression following TMZ. Interestingly, gene expression revealed that Interleukin-8 (IL-8) is significantly upregulated post-TMZ therapy (Supplementary Table 1). A recent report has demonstrated that epigenetic plasticity is a major driver of adaptation to targeted therapies in GBM8. To investigate the epigenetic regulation of plasticity during TMZ therapy, we next performed genome-wide ChIP-Seq analysis of TMZ-treated PDX GBM43 cells for histone 3 lysine 27 (H3K27) acetylation (ac), a marker of open chromatin and active gene transcription, and H3K27 tri-methylation (me3), a maker of closed chromatin and transcriptional repression. TMZ significantly augments H3K27ac levels at the IL-8 loci, as measured by ChIP summit in a previously established astrocyte-specific enhancer region9. This alteration was not accompanied by changes in H3K27me3 status (Figure 1B, Chromosome 4: 74783222-74783418, fold enrichment 3.02 as compared to input, p-value < 0.0001, FDR=0.004). Clearly, TMZ promotes a GIC state and alters the epigenomic landscape of GBM. Quantitative polymerase chain reaction (qRT-PCR) confirmed that TMZ treatment increased IL-8 mRNA levels in a time-dependent manner (Figure 1C) (***P < 0.0001). To investigate if this increase translates to higher protein levels and is maintained in vivo, immunofluorescence analysis was performed on a previously established orthotropic recurrent GBM model2 and confirmed that recurrent tumors had increased IL-8 expression (Figure 1E). In light of the ability of therapy to induce both IL-8 expression and a GIC state, we next examined if alterations in cell state can influence such expression without therapy. Freshly isolated PDX lines were cultured in GIC maintenance media (neurobasal media supplement with appropriate growth factors) or differentiation condition media (1% fetal bovine serum) with or without TMZ (50μM). Even without any chemotherapy, culturing GBM PDX lines in the GIC maintenance media significantly elevated the expression of IL-8 expression measured by enzyme-linked immunosorbent assay (ELISA, Fig 1C) (****P < 0.0001). Proneural subtype GBM43 and the classical subtype GBM6 expressed about 20 and 200 folds higher IL8, respectively, in the GIC maintenance media as compared to differentiation condition media. TMZ exposure induced IL8 expression in both culture conditions (****P < 0.0001). Moreover, this induction was specific to TMZ, as another anti-glioma alkylating agent BCNU failed to promote IL8 expression in any GBM lines tested (***P < 0.001 and ****P < 0.0001, Supplementary Fig. S1). Based on this observation, we set out to investigate the role of IL-8 in promoting therapy-induced cellular plasticity and disease recurrence. Critically, we utilized differentiation condition media (1% FBS or Neural Basal Media with BMP2), as these conditions initiate differentiation of GBM cells and enable us to observe how stimuli induce dedifferentiation to the GIC state during therapy [5, 11]. Culturing cells in a GIC-promoting media (Neurobasal supplemented with growth factors like EGF and FGF) would be inappropriate for this study, as it would force the cells to a GIC state and mask any GIC-inducing effects of our experimental manipulations.

**Figure 1:**
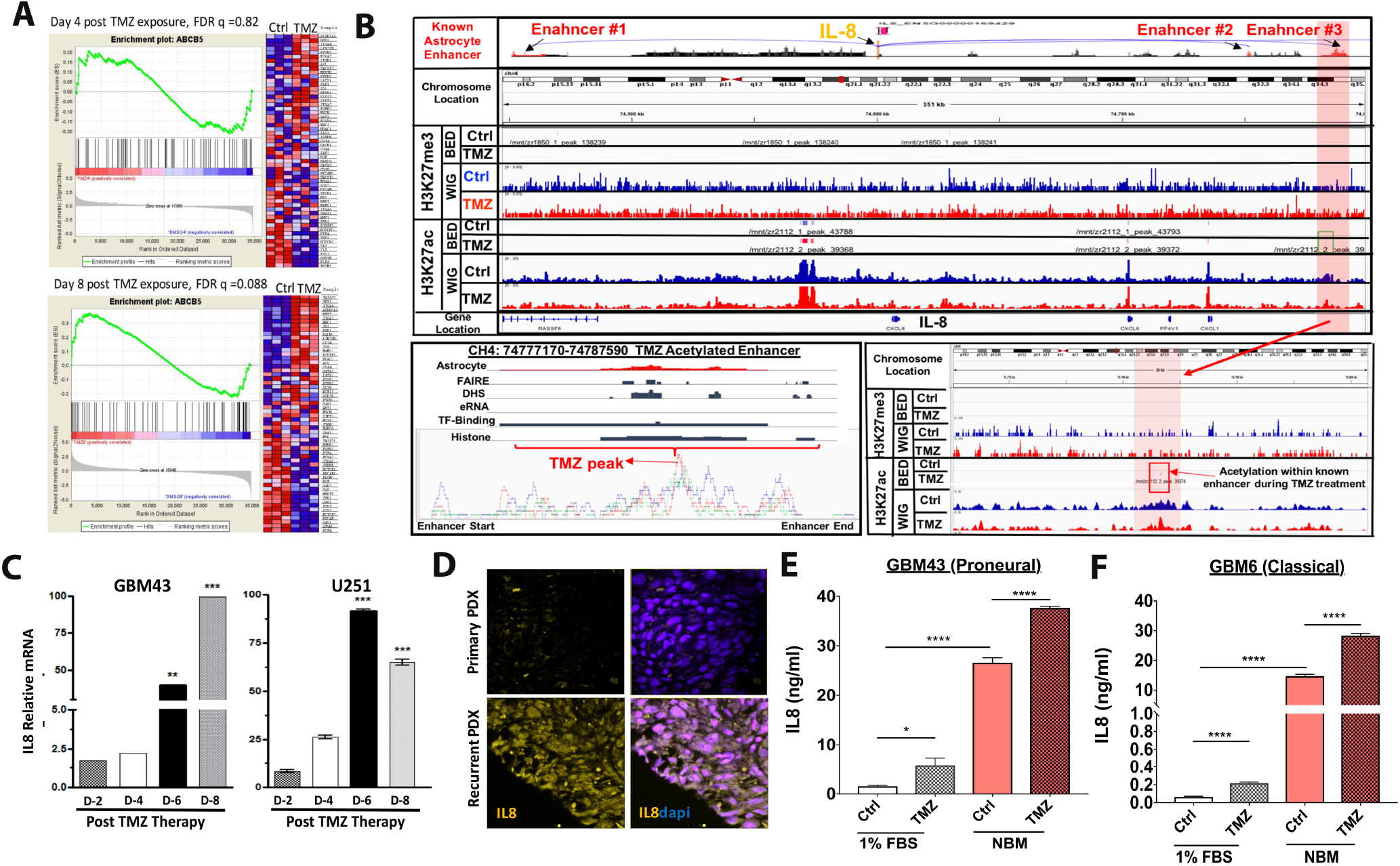
Therapeutic stress increases IL-8 expression. ***A***) PDX GBM43 cells were treated with TMZ (50μM) and RNA collected after four or eight days. Microarray analysis using Affymetrix platform was performed. Gene Set Enrichment Analysis (GSEA) for genes known to support and maintain the GIC phenotype [10] and demonstrated significance increase after 8 days [Day 4 FDR q 0.82 and Day 8 FDR .08 and FWER p value .046, TMZ-treated cells as compared to DMSO controls]. ***B***) Whole genome ChIP-seq analysis of H3K27 trimethylation (H3K27me) and H3k27 acetylation mark (H2K27ac) on PDX GBM43 following four days treatment with TMZ or vehicle control. Top two tracks represent the BET file showing significant enrichment of histone mark relative to DMSO control. Bottom two lanes represent WIG file showing significant enrichment peak. Analysis of three established astrocyte enhancer tracks for the IL-8 gene showed no changes in H3K27me3; however, they did show significant change in #3 enhancer region [Chr.4:74783222-74783418, fold enrichment 3.2 relative to IgG input. P<.0001. FDR=.004]. Right bottom inset image shows zoomed in view of the #3 enhancer region. Left bottom inset image represents overlayed peaks for TMZ and DMSO control. ***C***) Expression of IL-8 mRNA after exposure to 50μM TMZ was determined by quantitative real-time polymerase chain reaction (q-PCR) after treatment with TMZ across 8 days. All IL-8 values were normalized to glyceraldehyde 3-phosphate dehydrogenase (GAPDH). **Bars** represent means from three independent experiments and **error bars** represent the standard deviation. Multiple student T-tests were performed. **P<.01, ***P<.001. ***D***) Immunohistochemistry was performed on mouse brains with intracranial xenografts of GBM43. 1.5X10^5^ GBM43 PDX cells were implanted to establish orthotropic xenograft tumors. Animals received 2.5mg/kg of either DMSO or TMZ for 5 consecutive days, beginning 7 days after tumor implantation. Animals were sacrificed 5 days after the cessation of treatment, and whole brains were extracted, flash-frozen, then sectioned (8μm) and analyzed by immunofluorescence. 4’-6-diamidino-2-phenylindole (DAPI) stained DNA (**blue**) in the nuclei and allophycpcyanin-conjugated secondary (**orange**) antibody was used against primary antibody for IL-8. Dotted lines represent the edge of the tumor based on cell density and morphology. ***E-F***) PDX GBM xenografts were harvested and immediately plated in either mild differentiation media (DMEM containing 1%FBS) or GIC maintenance media (Neurobasal supplemented with FGF and EGF). Cells were then treated with either DMSO or TMZ (50μM). After four days, IL-8 levels were determined by ELISA. **Bars** represent means from three independent experiments and **error bars** represent the standard deviation. Multiple student T-tests were performed. **P<.01, ***P<.001.

### In silico analysis establishes IL-8 importance in GBM progression and patient outcomes

In order to examine the contribution of IL-8 to GBM clinical progression, we employed the Cancer Genome Atlas (TCGA) patient gene expression dataset. Our analysis illustrates that IL-8 transcript expression is elevated in World Health Organization (WHO) Grade IV glioma (GBM) (Figure 2A, IL-8 transcript expression, Grade IV vs Grade II *P* < 0.0001; Grade IV vs Grade III, P < 0.0001). Next, GBM patients were stratified based on IL-8 mRNA expression using quartile (Q1, Q3) split points; the lowest IL-8 quartile exhibited significantly higher median survival (Figure 2B) (All GBM: IL-8-down 15.1 months, IL-8-up 12.6 months, hazard ratio (HR) [95% CI] = 0.71[0.54,0.93], log-rank p value = 0.0112). This association between IL-8 and survival was especially pronounced in patients with proneural GBM tumors (Figure 2D) (Proneural subtype: IL-8-down 20.7 months, IL-8-up 9.3 months, HR[95%CI] = 0.54[0.33,0.87], log-rank p value = 0.0109). Multivariable stepwise Cox proportional hazards model confirmed IL-8 as an independent prognostic survival factor in all GBM patients, specifically those with proneural tumors (Figures 2B and 2D, right panel) (HR [95% CI]: All GBM 1.07[1.01, 1.14], P=0.0467; Proneural subtype 1.19[1.04, 1.37], P=0.014). Finally, average recurrence time in GBM IL-8-down-regulated patients is substantially longer than in IL-8-up-regulated patients (IL-8-down 47.9 months, IL-8-up 15.5 months, hazard ratio (HR) = 0.53[0.34,0.84], log-rank P value = 0.006) (Figure 2E).

**Figure 2:**
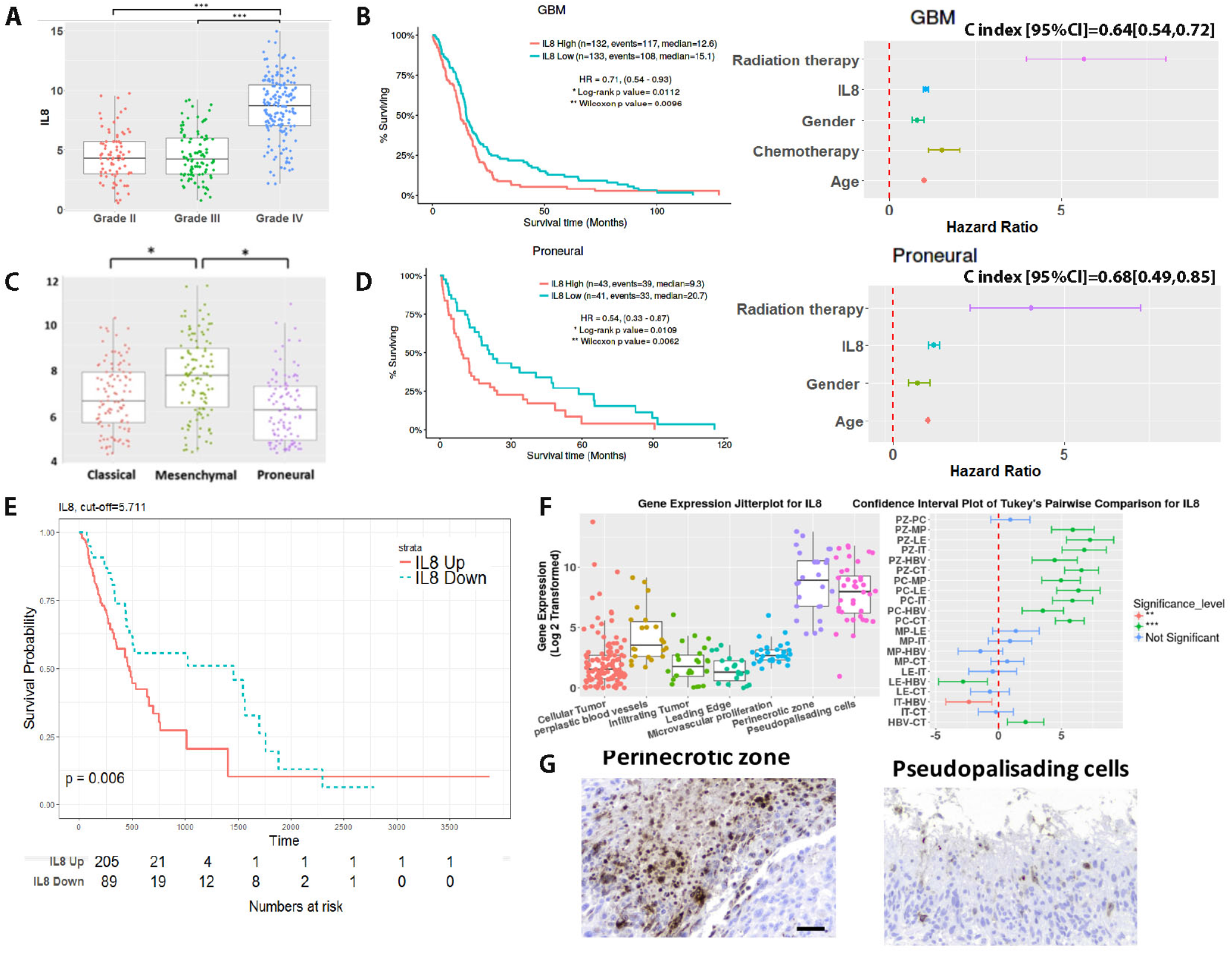
Bioinformatic analysis reveals IL-8 as a potential participant in Glioblastoma progression and outcomes. ***A***) *IL-8* mRNA expression levels were analyzed using the Affymetrix U133a platform on the Cancer Genome Atlas (TCGA) for different WHO grades of glioma. GBM (Grade IV glioma) patients had higher *IL-8* expression level than low-grade (Grade II, Grade III) glioma patients. ***B***) All GBM patients were stratified into IL-8-up-regulated and IL-8-down-regulated groups based on IL-8 gene expression using quartile (Q1, Q3) as split points. High expression of IL-8 correlated with reduced median survival. **Survival curves** were generated via the Kaplan-Meier method and compared by log-rank test. \*\**P*<.01. Multivariate stepwise Cox proportional hazards model with stepwise variable selection was conducted to examine whether IL-8 could be an independent factor for predicting survival with major clinical variables adjusted. This analysis confirmed that IL-8 was an independent prognostic factor for survival in all GBM patients (HR [95% CI]: All GBM 1.07[1.01, 1.14], P=0.0467). **C Index** (95%CI) or C statistics are provided. ***C***) Within Grade IV glioma subtypes, proneural GBM patients had the lowest level of IL-8 expression. **Box plots** represent means and interquartile range. One-way ANOVAs with Bonferroni correction for the multiple comparisons were performed. \**P*<.05, \*\*\**P*<.001. ***D***) Patients with proneural GBM were stratified into IL-8-up-regulated and IL-8-down-regulated groups based on IL-8 gene expression using quartile (Q1, Q3) as split points. Kaplan-Meier survival curves and multivariate stepwise Cox proportional hazards models were generated as in B. ***E***) In TCGA database (U133a) GBM patients with IL-8-down-regulated also have a longer time to recurrence, compared to IL-8-up-regulated patients. ***F***) The Ivy Glioblastoma Atlas Project (IVY GAP) was employed to determine the location of IL-8 in glioblastoma samples. Each column represents the data for one biopsy from a tumor. Microdissection for the noted anatomically portions of the tumor and subsequent mRNA extraction and expression analysis demonstrated that IL-8 is upregulated in the perinecrotic zone and pseudopalsading cells. **Heatmap** illustrates most significantly and differential expressed genes with a false discovery rate <0.01. mRNA expression in each anatomical compartment were compared. IL-8 was significantly upregulated in the pernecrotic zone. **Bars** represent means from three independent experiments and **error bars** represent the standard deviation. Multiple student T-tests were performed. ***P<.001. ***G***) Brain tumor samples from primary biopsies or surgical resections were stained for IL-8 at the Northwestern Brain Tumor Tissue Bank. Histological and morphological analysis confirm that IL-8 is present in the perinecrotic zone and pseduopalsading cells. **Scale bar** 50 microns.

Next, to investigate IL-8 expression patterns across different tumor compartments, we utilized the Ivy Glioblastoma Atlas Project (IVY GAP) [12], which demonstrated that IL-8 mRNA is elevated in pseudopalsading cells and the perinecrotic zone, two areas linked to the GIC subpopulation (Figure 2F). All of these data further justify our interest in IL-8 as a critical participant in GBM progression and therapy-induced plasticity.

### Immunohistochemical analysis of IL-8 expression in matched primary recurrent GBM tissue

To investigate IL-8 expression in GBM tissue from patient samples, 75 GBM specimens from Northwestern University’s Brain Tumor Bank were subjected to immunohistochemical characterization. Pathological analyses found approximately 65% (49/75) of GBM samples were IL-8-positive. Further, 17 matched primary and recurrent GBM tumor pairs were examined for IL-8 expression. We found elevated IL-8 in 65% (11/17) of recurrent tumors (Figure 3A, 3B, and 3C). Critically, IHC analysis revealed high IL-8 expression in the typically hypoxic pernecrotic zone, an anatomical compartment where niche for the GIC was reported to be maintained 11-14 (Figure 2F). Our analysis also revealed that both tumor cells and infiltrating macrophages express IL8 (Figure 3D and E).

**Figure 3:**
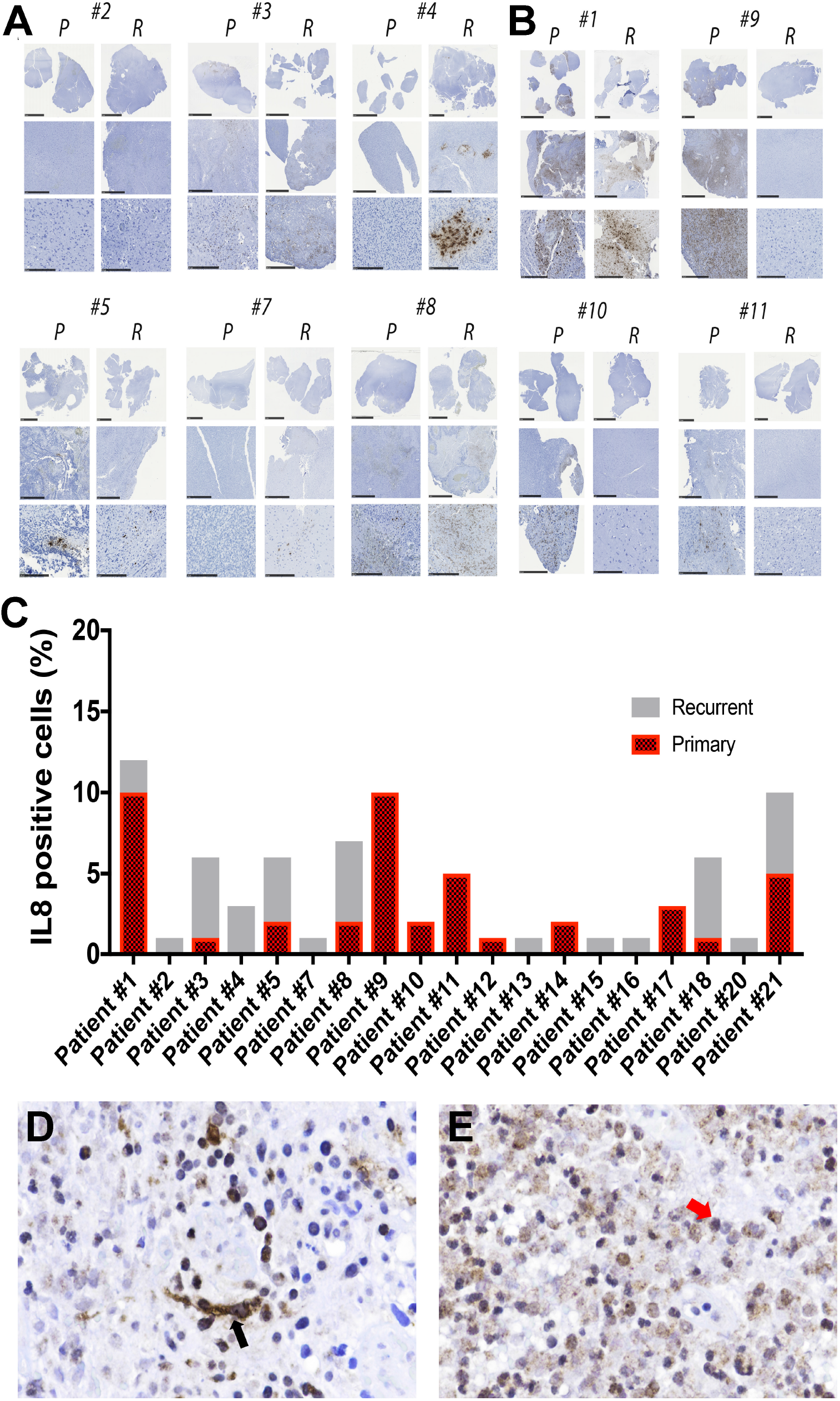
Immunohistochemical analysis of IL-8 expression in the matched primary and recurrent GBM patient samples. ***A***) Representative immunohistochemical staining for IL-8 in the matched primary (P) and recurrent (R) clinical GBM samples. Top row low-power and bottom row high-powered magnification of IL-8 staining. **Scale bar** for all top row images 1mm, all middle row images 5mm and for the bottom row images 250μm. These sets of patient samples show IL-8 upregulation in the matched recurrent tissues. ***B***) Same as previous, but this set of patient samples shows a decrease in IL-8 staining for recurrent GBM as compared to their matched primary GBM. ***C***) Quantitative analysis of the percent of IL-8 positive cells in the matched primary and recurrent GBM tissue performed by our in-house neuropathologist. Representative Immunohistochemical analysis of IL-8 expressing ***D***) tumor cells (left, black arrow) and ***E***) infiltrative macrophage (right, red arrow).

### IL-8 receptor CXCR2^+^ GBM cells acquire CD133 expression during anti-glioma chemotherapy

Next, we set to investigate how IL-8 signaling influence GBM proliferation and signaling. CXC motif chemokine receptors 1 and 2 (CXCR1 and CXCR2) are the major receptors for IL-8 [13]. To investigate their role in IL-8 mediated signaling in GBM, we first interrogated TCGA data. We observed that expression of both these receptors was significantly elevated in GBM tumors compared to low grade gliomas (Figure S2-A). Analysis of our ChIP-seq data showed post therapy (4 days following exposure to 50μM TMZ) accumulation of open chromatim mark at a known enhancer site[14] for CXCR2 (Figure 4A, Chromosome 2: 218714857-218715098, fold enrichment 3.15 as compared to IGG input, p value < 0.0001, FDR=0.0004; Chromosome 2: 218714701-218715149, fold enrichment 3.63 as compared to IGG input, P value < 0.0001, FDR< 0.0001), as well as significant decreases in H3K27me3 levels in the gene body (Figure S2 C-H, P value < 0.001). In contrast, gene body H3K27me3 levels were significantly increased in the CXCR1 gene loci (Figure S2-B). Next, CXCR1/2 expression post-TMZ treatment was examined by FACS analysis. All cell lines increased expression of both receptors posttreatment, with the most pronounced differences in CXCR2 (Figure 4B, P = 0.00015). Moreover, CXCR2 expression was also elevated in the CD133+ GIC population (Figure 4C). Time course FACS following TMZ treatment revealed that a CXCR2^+^ cell population exists prior to TMZ treatment; this population rapidly gains CD133 expression during treatment (Figure 4D). CXCR1 expression was not altered during therapy (Figure 4B and Figure S2-I). Analysis of the downstream effectors of the IL-8/CXCR signaling cascade in PDX GBM demonstrated that IL-8 induces dose-dependent activation of canonical ERK1/2 and AKT signaling (Figure S2-J)

**Figure 4:**
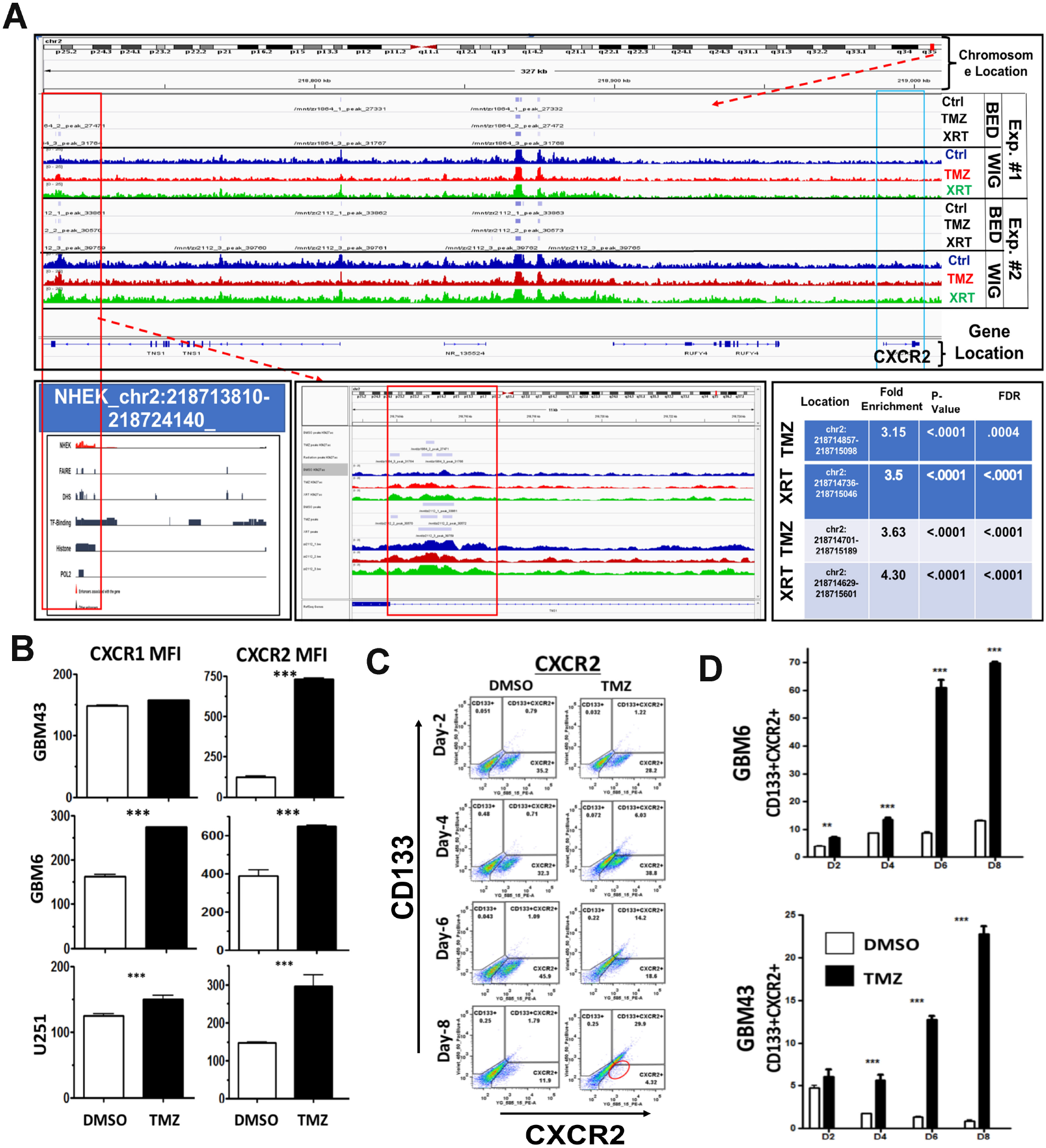
Therapeutic stress alters epigenetic status and increases expression of CXCR2, one of the major receptors for IL-8. ***A***) ChIP-seq analysis was performed for H3K27ac, a marker of open chromatin associated with activation of gene expression, and H3K27me, associated with closed chromatin and repressed gene expression, on PDX GBM43 cells. Cells were treated with either TMZ (50μM) or equimolar DMSO for four days prior to analysis. Top track shows the location of the CXCR2 gene, with higher magnification analysis of the area shown below. TMZ-treatment led to increased H3K27ac enrichment, including in a well-established enhancer region for CXCR2 (Green box and red box) [Chr2:218714857-218715098, fold enrichment 3.14 relative to IgG input, p value <0.0001, FDR 0.0004; Chr2:218714701-218715149, fold enrichment 3.63 comp. input, p<0.0001, FDR<0.0001]. Further, gene body H3K27me3 were significantly reduced following TMZ [P=00120 in TMZ relative to DMSO control levels]. ***B***) FACS analyses were performed to determine how TMZ treatment alters the levels of CXCR2 in three GBM cell lines— GBM43, GBM6, and U251. Samples were analyzed 8 days after initial treatment with either DMSO or TMZ (50μM). All data is expressed as the mean fluorescent intensity (MFI). **Bars** represent means from three independent experiments and **error bars** represent the standard deviation. Multiple student T-tests were performed. \*\*\**P*<.001. ***C***) Representative FACS scattered plot analyses from GBM43 cells treated with either DMSO or 50μM TMZ across 8 days. **Circle** highlights the clear shift of the CXCR2 expressing population into the CD133+ compartment. (D) TMZ treatment significantly increased expression of CXCR2 in both GBM43 and GBM6, with CXCR2 expressing cells beginning to co-express CD133 GIC markers in a time dependent manner. All data are expressed as the percentage of total live cells. **Bars** represent means from three independent experiments and **error bars** represent the standard deviation. Multiple student T-tests were performed. \*\**P*<.01; \*\*\**P*<.001.

### IL-8 increases the self-renewing capacity of GBM cells and the expression of GIC markers

To determine if the IL-8-CXCR signaling axis promotes induction of the GIC state in GBM, we performed extreme limiting dilution assay (ELDA) on GBM6 and GBM43 PDX cells in neurosphere media containing IL-8 (50 ng/ml). IL-8 increased GIC frequency about 3.3-fold for GBM43 and 2.3-fold for GBM6 (Figure 5A, P=0.001). We next examined how IL-8 alters the expression of known GIC promoting genes, using our proprietary GIC-specific reporter cell line[11]. IL-8 activation significantly increased reporter activity (Figure 5B and Figure S3-A for SOX2-RFP reporter and Nanog-RFP reporter P = 0.0005). Next, to investigate how IL-8-CXCR signaling cascade interacts with the network of GIC-promoting genes in patient tumors, we selected the top GIC-associated genes activated during TMZ therapy (Figure 1A and Supplementary Table 2) and correlated with their levels with IL-8 mRNA using the GlioVis data portal 16-18. IL-8 expression was significantly correlated with critical GIC associated genes including KLF4, CD44, HIF1A, HIF2A, Myc and Twist expression (Figure 5C). To examine the IL-8 expression in different anatomical location and potential colocalization of these GIC-specific genes with areas of high IL-8 transcript level, we employed the Ivy Glioblastoma Atlas project; we observed that these genes transcripts were correlated with the IL-8 expression in both the perinecrotic zone and pseudopalsading cells (Figure 5C, heat map; Ivy Glioblastoma Atlas Project. Available from http://glioblastoma.alleninstitute.org). Immunoblot analysis of PDX lines exposed to IL-8 show time-dependent induction these genes (Figure 5D) as well as various critical GIC-associated transcription factor such as C-myc, Nanog, Sox2, OCT4 (and Figure S3-D). Finally, to examine the role of post-therapy IL-8 in inducing the GIC specific gene expression, we combined DMSO or 50μM TMZ with either control IgG or anti-IL-8 antibody. Blocking IL-8 both reduced basal expression of GIC markers and prevented TMZ-induced increases in SOX2 and C-Myc (Figure 5E), indicating IL-8 participates in activating genes capable of promoting GIC niche during therapeutic stress.

**Figure 5:**
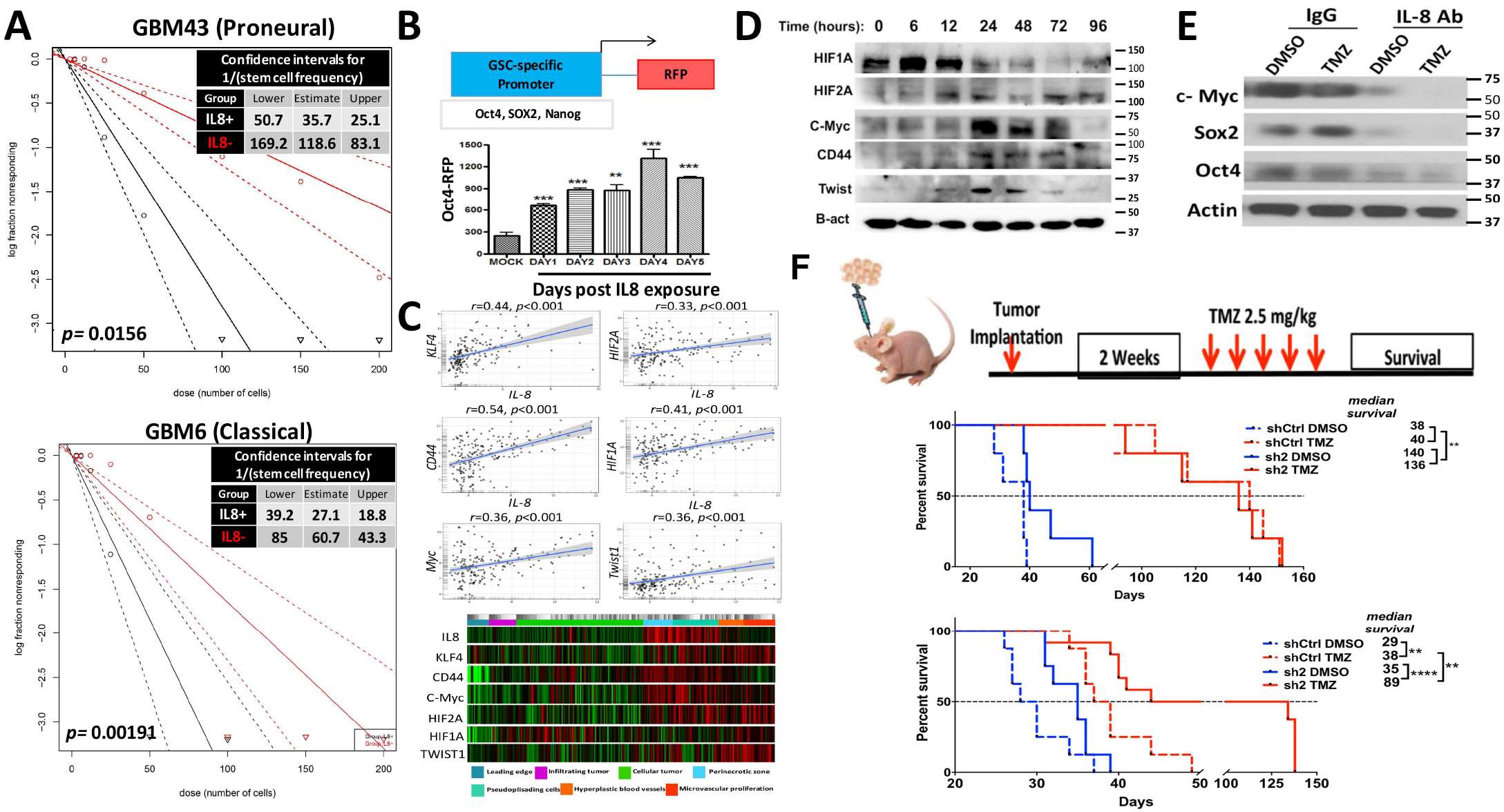
IL-8 contributes to glioma-initiating cell phenotype and contributes to GBM growth in vivo. ***A***) Limiting dilution neurosphere assays were performed on two cell lines— GBM43 and GBM6—after treatment with 50 ng/ml of IL-8. Stem cell frequency for GBM43 with IL-8 35.7, lower limit 50.7 and upper limit 25.1 as compared to no IL-8 118.6, lower limit 169.2 and upper limit 83.1, p=0.0156; for GBM6 with IL-8 27.1, lower limit 39.2 and upper limit 18.8 as compared to no IL-8 60.7, lower limit 85 and upper limit 18.8, p=0.001. ***B***) To determine the ability of IL-8 to influence cellular plasticity, we employed a reporter cell line in which RFP expression is controlled the OCT4 promoter. Cells were treated with 50 ng/ml of IL-8 and RFP expression was monitored by FACS over 6 days. Treatment increased both Oct4 and Sox2. **Bars** represent means from three independent experiments and **error bars** represent the standard deviation. Multiple student T-tests were performed. **P<.01, ***P<.001. ***C***) The network of GIC-promoting genes in patient tumors, we selected the top GIC-associated genes activated during TMZ therapy (Fig. 1A and Supplementary Table 2) and correlated with their levels with IL-8 mRNA using the Ivy Glioblastoma Atlas Project. Available from (http://glioblastoma.alleninstitute.org) data portal 16-18. IL-8 expression was significantly correlated with critical GIC associated genes including KLF4, CD44, HIF1A, HIF2A, Myc and Twist expression (Fig. 5C). The IL-8 expression in different anatomical location and potential colocalization of these GIC-specific genes with areas of high IL-8 transcript level (Figure 5C, heat map). ***D***) Immunoblot analysis of endogenous glioma-initiating cell-associated transcription factors expression upon stimulation with escalation dose of IL-8 (0-100 ng/ml) for 24 hours. Protein extract of IL-8 treated PDX line GBM43 and GBM6 were immunoblotted with antibody against several GIC markers, including c-myc, Sox2, Nanog, KLF4, OCT4 or an antibody against ß-actin as a control for equal loading. ***E***) GBM43 PDX cells were treated with neutralizing antibody against IL-8 or control IgG antibody (100 ng/ml) prior to treatment with DMSO or 50μM TMZ. Neutralizing antibody was added every day for 8 days and protein extracts from this experiment were immunoblotted with antibody against c-myc, Sox2, OCT4 or an antibody against ß-actin as a control for equal loading. ***F***) Schematic diagram of experiment design for *in vivo* testing. Top graph, U251 cells were infect with lentivirus (Sigma Mission shRNA) shRNA against IL-8 or scrambled shRNA (control) with 10 infectious unit/cell. 2X10^5^ cells transduced cells were stereotactically injected into the right hemisphere of the brain of athymic nude mice (n=8 per group, 4 male and 4 female). 2 weeks after implantation, two groups of mice, control and knockdown, were treated with vehicle treated (DMSO, top curve) or TMZ (2.5 mg/kg) intraperitoneally. Survival curves were obtained by the Kaplan-Meier method, and overall survival time was compared between groups using log-rank test. All statistical tests were two-sided. Bottom graph, to examine the role of IL-8 in GBM progression in a more clinically relevant manner, next the same method was used to knock down the IL-8 expression in GBM43 PDX line. 1.5X105 cells were injected stereotactically injected into the right hemisphere of the brain of athymic nude mice (n=8 per group, four male and four female). Survival curves were obtained by the Kaplan-Meier method, and overall survival time was compared between groups using log-rank test.

### IL-8 enhances GBM growth and therapy resistance in vivo

To elucidate IL-8’s role during *in vivo* GBM growth, U251 cells with stable IL-8 knockdown were established using short hairpin RNA (shRNA) technology. IL-8 secretion was effectively knocked down by two shRNA constructs (Figure S4-A and B, P<0,0005). Proliferation capacity was altered significantly in cells with the highest IL-8 knockdown compared to control population (Figure S4-C, P=0.018), but cell cycle profiles remained unchanged across all populations (Figure S4-D & E). Athymic, immunodeficient mice then had GBM cells expressing either sh-Control or anti-IL-8 shRNA#1 implanted into the right cerebral hemisphere. Each group was then divided into two groups that received either DMSO or TMZ (2.5 mg/kg, i.p)(n=7/group). Treatments were initiated two weeks postimplantation for five consecutive days. In this model, IL-8 knockdown significantly increased the median survival of animals with orthotropic GBM regardless of chemotherapy exposure (Figure 5F, first graph median survival sh-control 38 days vs. sh#1 IL-8 140 days; hazard ratio of survival = 4.737, 95% CI = 4.190 to 101.3, P=0.0021). In a clinically relevant model, PDX GBM43 (which express high basal levels of IL-8, Figure 1C & E) were infected with a lentivirus (10 infectious unit/cell) carrying shRNA against IL-8, reducing IL-8 expression by 50% (Figure S6-A, P>0.0005) after transient transfection. Implantation of these IL-8 knockdown GBM43 prolonged median survival about 38% compared to control shRNA (Figure 5F, 2^nd^ graph median survival for sh-control 29 days vs. sh#2 IL-8 40 days P=0.0003). Moreover, IL-8 knockdown significantly enhanced the therapeutic efficacy of TMZ and improved survival about 51 days (Figure 5F).

### IL-8 signaling promotes epigenetic alterations in GBM

To identify potential mechanisms by which IL-8 influences growth and promotes therapy resistance, we analyzed correlations between IL-8 and 12042 other genes using TCGA patient data via Pearson correlation coefficients. Our analysis returned 68 genes with coefficients >0.5 or <-0.5 and false discovery rate (FDR) <0.05 that correlate with IL-8; we then conducted unsupervised hierarchical clustering. Observed cophenetic correlation determined optimal clusters, which we validated by silhouette plots and principal component analysis (PCA). Two clusters with the highest cophenetic coefficient at 0.95 and average silhouette width at 0.46 (Figure S5 and Figure 6A) separated well upon visualization of PCA. Using the Enrichr platform, we determined that the first of these clusters (Group A) is involved in regulating cell chemotaxis (GO: 0060326, Adjusted P-value 1.058e-11) and cytokine activity (GO: 0005125, Adjusted P-value 2.495e-9), well-established canonical roles for IL-8[15]. Cluster B includes genes enriched for wounding (GO: 009611, Adjusted P-value 0.002) and hypoxia (GO: 0001666, Adjusted P-value 0.004), also known IL-8 connections [16, 17]. Interestingly, IL-8 signaling also positively correlated with genes known to regulate epigenetic processes, specifically histone3 lysine27 trimethylation (H3K27me3) (GO: 0001666, Adjusted P-value 0.03).

**Figure 6:**
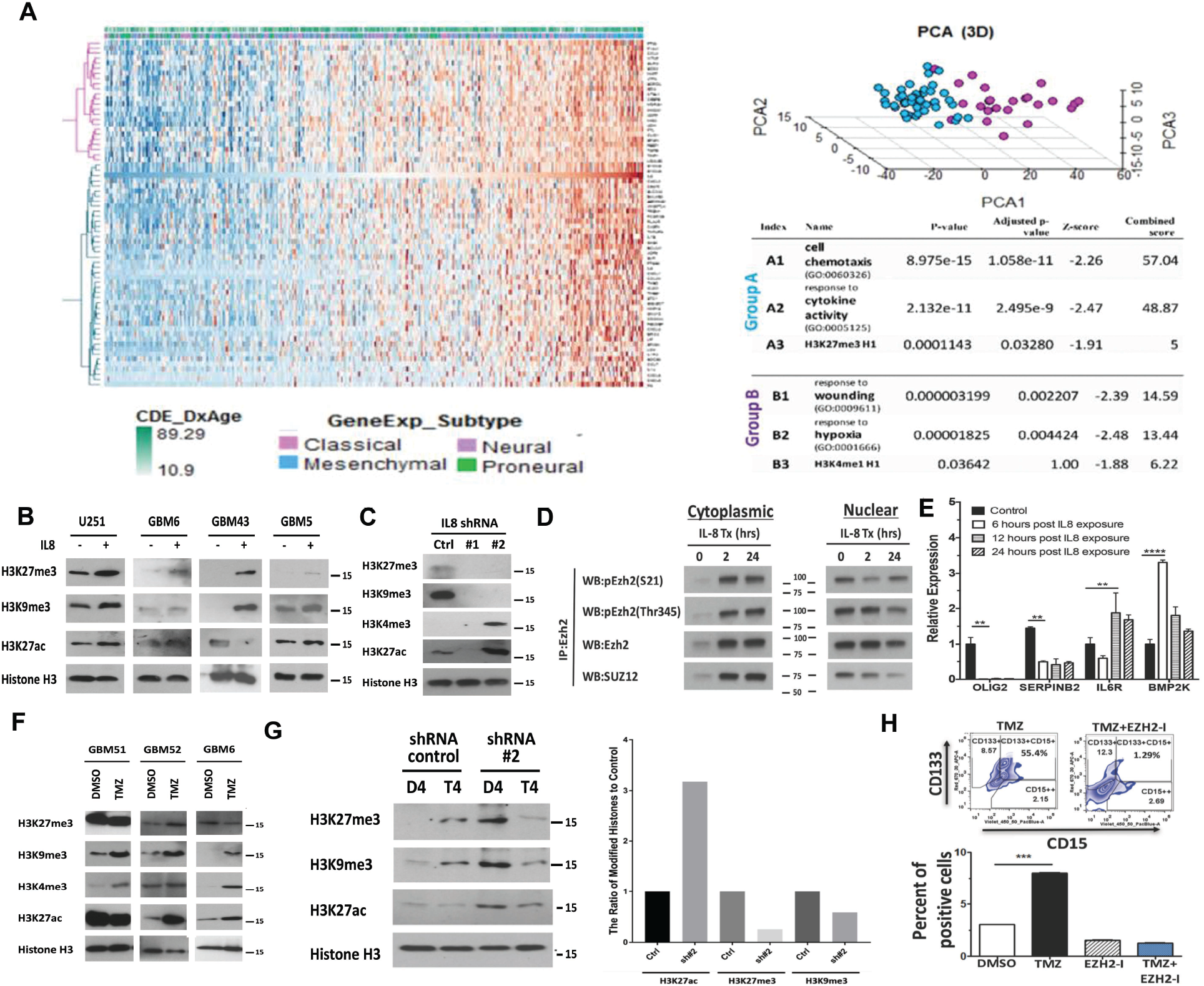
IL-8 signaling alters histone marks, promoting post-therapy epigenetic plasticity. ***A***) Correlation between IL-8 and 12042 genes from TCGA was determined by Pearson correlation coefficients. 68 genes with coefficients >0.5 or <-0.5 and false discovery rate (FDR) <0.05 were selected. Unsupervised hierarchical clustering of those genes found two clusters with the highest cophenetic coefficient at 0.95 and an average silhouette width at 0.46. Principal component analysis was used to validate these clusters. Enrichment analysis found one cluster of genes was enriched at cell chemotaxis (GO: 0060326, Adjusted P-value 1.058e-11) and cytokine activity (GO: 0005125, Adjusted P-value 2.495e-9), while another cluster of genes enriched at wounding (GO: 009611, Adjusted P-value 0.002) and hypoxia (GO: 0001666, Adjusted P-value 0.004). ***B***) Representative immunoblot of different histone marks. A panel of PDX lines from a different subtype of GBMs was exposed to IL-8 (50ng/ml) for 24 hours. Nuclei were extracted from the harvested cells and subjected to immunoblot analysis for suppressive histone marks H3K27 and H3K9 trimethylation (me3) and activating mark H3K27 acetylation(ac). Immunoblotting for total histone three was performed to confirm the equal loading. ***C***) The extracted nuclei from the U251 IL-8 knockdown cells as described in Figure 5F were subjected to immunoblot analysis for various histone marks as described above. ***D***) GBM43 cells were treated with IL-8 (50ng/ml) for 2 and 24 hours, cytoplasmic and nuclear extract was prepared, cells were harvested, and immunoprecipitation assays were performed with the anti-EZH2 antibody. Immunoprecipitated protein was subjected to immunoblot analysis with antibodies against phosphor-EZH2 S21 and Thr345 and SUZ12. ***E***) IL-8 (50 ng/ml)-treated GBM43 PDX lines were harvested at 6, 12 and 24 hours post IL-8 exposure. mRNA was extracted and subjected to reverse-transcription polymerase chain reaction (RT-PCR) analysis of *OLIG2, SERPINB2, IL6R and BMP2K* transcripts. Bars represent means from two experiments in triplicate and error bars represent the standard deviation. Multiple student T-tests were performed. **P<.01, ****P< 0001. ***F***) A panel of GBM PDX lines was treated with TMZ (50μM) for 48 hours. Nuclei were extracted from the harvested cells and subjected to immunoblot analysis for suppressive histone marks H3K27me3 and H3K9me2 and activating mark H3K27ac and H3K4me3. Immunoblotting for total histone three was performed to confirm equal loading. ***G***) The U251 IL-8 control and knockdown cells as described in figure 5A were treated with TMZ(50μM) for four days. Nuclei were extracted from the harvested cells and subjected to immunoblot analysis for suppressive histone marks H3K27me3 and H3K9me2 and activating mark H3K27ac. Left, representative densitometry analysis is expressed as percent of control shRNA (shCtrl). ***H***) The GBM43 PDX line was treated with TMZ (50μM) in the presence of 3-Deazaneplanocin A (DZNep, EZH2-I, 5 μmol/L), a histone methyltransferase Ezh2 inhibitor for 8 days. Cells were harvested and the GIC population was analyzed by FACS analysis of the CD133 and CD15 positive cells. Bars represent means from two experiments in triplicate and error bars represent the standard deviation. Multiple student T-tests were performed. ***P<.001.

Trimethylation of H3K27 suppresses gene expression via recruitment of the Polycomb repressor complex (PRC), predominately regulated by two methyltransferases, EZH2 and G9a [18, 19]. Confirming our in silico result, treatment with IL-8 significantly increased trimethylation of H3K27, as well as another PRC complex target, H3K9[20], in three PDX lines (Figure 6B). Moreover, reduction of IL-8 expression via shRNA abolished methylation of the H3K27 and H3K9 residues (Figure 6C), with dose-dependent effects (Supplementary Figure S6-B). IL-8 signaling alters histone status in GBM cells.

To further elucidate the connection between IL-8 and PRC, we next examined the status of crucial PCR members following IL-8 exposure. Phosphorylation of EZH2 in response to various extracellular stimuli remodels the epigenomic landscape, allowing cellular adaptation [21]. Specifically, phosphorylation at Thr345 enhances recognition of target genes leading to recruitment of PRC2 and suppression of transcription via H3K27me3[22]. Contrastingly, extracellular signaling like AKT suppresses the methyltransferase activity of Ezh2 by phosphorylating Ser21[23]. Our previous results illustrated that IL-8-CXCR interaction could activate various downstream signaling cascades, including PI3K-AKT (Figure S6-C & D)[24]. We, therefore, examined the alteration of the phosphorylation status of Ezh2 by IL-8 via immunoprecipitation (IP) in the nuclear and cytoplasmic fraction (Figure 6D). IL-8 stimulation enhanced phosphorylation of Ezh2 at both Ser21 and Thr345, exclusively in the cytoplasmic fraction. However, within two hours of IL-8 stimulation, S21-phosphorylated (inhibited) EZH2 levels were decreased in the nucleus, while Thr345-phosphorylated (activated) Ezh2 accumulated. Binding of EZH2 to SUZ12, an essential PRC protein, increased only in the cytoplasmic compartment after IL-8 exposure, while the nuclear accumulation of Ezh2-SUZ12 complex gradually decreased. We conclude that within two hours of IL-8 exposure, the PRC2 activity may increase, but by 24 hours nuclear accumulation of PRC2 complex is reduced by heightened phosphorylation of EZH2 at Ser21.

To determine the functional effect of IL-8 on EZH2 activity, we analyzed two genes positively regulated by Ezh2 (OLIG2, SERPINB2) and two negatively regulated genes IL6R and BMP2K via qRT-PCR (Figure 6E)[25]. Remarkably, IL-8 exposure reversed expression of all four Ezh2 target genes, indicating that IL-8-induced Ezh2 modifications do in fact alter gene transcription.

Next, we expanded our investigation into TMZ-induced therapeutic stress-induced alterations in histone markers that are targets of Ezh2/PRC2. Treatment with TMZ induced subtype-and time-dependent global changes in epigenetic markers H3K27me3 and H3K9me3 and open chromatin marker H3K27 acetylation (ac) (Figure 6F). Additionally, PRC2 target H3K9me3 was upregulated in GBM6. H3K27 trimethylation and acetylation were upregulated within 48 hours post-TMZ exposure and stayed elevated. H3K4 trimethylation also increased within 96 hours, indicating acquisition of bivalency[26].

Given that TZM induces IL-8 signaling and Ezh2-dependent changes in histone status, we next examined the relationship between TMZ, IL-8, and Ezh-2 target histones. We treated IL-8-knockdown cells with TMZ for 96 hours and analyzed histone status. Reduced IL-8 levels abolished methylation of Ezh2 targets H3K27 and H3K9; however, H3K27ac decrease was minimal. Consequently, we suspect that chemotherapy-induced IL-8 signaling participates in for Ezh2-dependent epigenetic modifications during therapeutic stress. Finally, to investigate the role Ezh2/PRC2 complex activity in promoting therapy-induced cellular plasticity the GBM43 PDX line was treated with TMZ in the presence of 3-Deazaneplanocin A (DZNep, EZH2-I), a histone methyltransferase Ezh2 inhibitor. As shown in Figure 6H, DZNep completely abolishes the induction of a GIC population after therapy, as measured by FACS analysis of the CD133+ and CD15+ populations (Figure 6H, P>0.0005).

## Discussion

The ability of GBM cells to adapt to current therapies and generate treatment-resistant recurrences represents one of the critical challenges facing brain tumor researchers and clinicians. Here, we provide data that sheds new light on mechanisms that may underlie this powerful ability to react to and overcome standard of care therapies. This study highlights the IL-8/CXCR2 signaling pathway as a critical player in this process and a potential target for blocking GBM cellular plasticity during therapy. Specifically, our data show: (1) therapeutic stress alters the epigenetic regulation of IL-8 leading to increased expression and secretion of IL-8; (2) bioinformatics analysis and IHC analysis of matched primary and recurrent patient tissues suggest that IL-8 significantly influences patient progression and time to recurrence; (3) therapeutic stress induced IL-8 alters the phenotype of GBM cells, shifting them to a more GIC-like state; (4) IL-8 supports GBM aggression and resistance to chemotherapy in vivo; and (5) IL-8 signaling may influence the acquisition of GIC state via modulation of the histone modifying PRC2 complex. Overall, these results provide key insights into the role of IL-8 in promoting the formation of aggressive, treatment-resistant recurrent tumors.

Induction of cellular plasticity, primarily when it promotes the adoption of therapy by inducing a GIC phenotype, is a well-established player in the formation of GBM recurrence. Indeed, several other groups and we have demonstrated how standard therapies can initiate this process and enrich GBM tumors with GIC cells [4, 5, 11, 27]. The ability to block this cellular plasticity remains a critical goal of contemporary research into novel therapies for GBM. However, targetable players in this process have yet to be identified. Here, we provide some convincing evidence that IL-8 represents one such target. Our results show that IL-8 is sufficient to induce the GIC state on its own. Further, we show that it is both adequate and necessary for the adoption of GIC state during therapeutic stress. These results indicate that IL-8 signaling is a potent regulator of GBM phenotype. Not only was IL-8 able to induce the expression of CD133 and CD15, two GIC phenotypic markers, it also caused increased expression of many transcription factors known to promote the GIC phenotype and recurrence, including HIF, c-myc, Sox-2, and CD44. In light of the ongoing debate regarding the precise gene expression profile of GICs, this robust induction, combined with our matched patient data, provides strong evidence that IL-8 is capable of activating cell reprogramming towards a more GIC-like state during anti-glioma chemotherapy.

Another key aspect of this study concerns the tumor microenvironment and how anticancer therapeutic can influence its composition. As previously mentioned, environmental cues are key regulators of GBM plasticity and the generation of recurrent growth. Here, we show that GBM cells actively sculpt their microenvironment following treatment with chemotherapy. This fact further corroborates a growing body of evidence that tumor cells actively cultivate a pro-growth microenvironment. An interesting facet of our data is the fact that culturing GBM cells in pro-GIC conditions alone led to an increase in IL-8 levels, suggesting that tumor cells may utilize positive-feedback loops to respond to their surroundings rapidly. These positive feed-forward loops may represent prime targets for combination therapy with traditional anti-tumor treatments.

One open question from this study is potential non-tumor sources of IL-8 in the tumor microenvironment. While our results illustrate a robust role for GBM autocrine IL-8, it remains an open question how IL-8 from surrounding non-tumor cells might participate in the recurrence process. Cytokines from the stroma and infiltrating immune cells have been identified as regulators of tumor behavior in other cancer types, including breast cancer [28]. In fact, our IHC analysis shows that infiltrating macrophage express IL-8 in the tumor microenvironment. Moreover, astrocytes and brain endothelial cells are both known to release cytokines, especially in response to damage [29, 30]. Indeed, one of the top hits from our bioinformatics analysis of pathways correlated with IL-8 expression in patient samples was wound healing. These data and the potential involvement of healthy nontumor cells provide further evidence for the theory that cancer represents an “un-healing” wound and is caused by inappropriate activation of damage response pathways. New research will shed light on this hypothesis. In light of the fact that GBM is highly sensitive to changes in the microenvironment [2, 31] and factors secreted by nearby neurons[32], it is highly possible that non-tumor IL-8 may participate in the processes described here. This question, however, does not change the fact that IL-8 represents a potentially druggable target in GBM.

Our data indicate for the first time that IL-8 is capable of causing alterations in the epigenetic status of key gene regulation factors, such as H3K27. While evidence increasingly shows the importance of epigenetic regulation in GBM growth and therapy resistance8, the mechanisms activating these processes remain incompletely understood. Our work indicates that IL-8 represents one tumor-derived trigger for activating epigenetic responses to tumor therapy via modulation of the canonical PRC2 complex.

In sum, our data show that IL-8 is a key microenvironmental factor involved in promoting cellular plasticity in GBM. Through analysis of murine models and patient data, we illustrate the high degree of influence IL-8 holds over tumor progression. Further, our work connects IL-8 signaling to the increasingly important area of epigenetic regulation of gene expression in the service of tumor growth. These results highlight IL-8/CXCR signaling as a key target for GBM drug development, especially in combination with standard of care therapies.

## Acknowledgement

We like to thank Ting Xiao for her data analysis assistant.

